# Genome-wide association studies suggest limited immune gene enrichment in schizophrenia compared to five autoimmune diseases

**DOI:** 10.1101/030411

**Authors:** Jennie G. Pouget, Vanessa F. Gonçalves, Schizophrenia Working Group of the Psychiatric Genomics Consortium, Sarah L. Spain, Hilary K. Finucane, Soumya Raychaudhuri, James L. Kennedy, Jo Knight

**Author notes:** Address correspondence to Jennie Pouget.

## Abstract

There has been intense debate over the immunological basis of schizophrenia, and the potential utility of adjunct immunotherapies. The major histocompatibility complex is consistently the most powerful region of association in genome-wide association studies (GWASs) of schizophrenia, and has been interpreted as strong genetic evidence supporting the immune hypothesis. However, global pathway analyses provide inconsistent evidence of immune involvement in schizophrenia, and it remains unclear whether genetic data support an immune etiology *per se*. Here we empirically test the hypothesis that variation in immune genes contributes to schizophrenia. We show that there is no enrichment of immune loci outside of the MHC region in the largest genetic study of schizophrenia conducted to date, in contrast to five diseases of known immune origin. Among 108 regions of the genome previously associated with schizophrenia, we identify six immune candidates (*DPP4, HSPD1, EGR1, CLU, ESAM, NFATC3*) encoding proteins with alternative, nonimmune roles in the brain. While our findings do not refute evidence that has accumulated in support of the immune hypothesis, they suggest that genetically mediated alterations in immune function may not play a major role in schizophrenia susceptibility. Instead, there may be a role for pleiotropic effects of a small number of immune genes that also regulate brain development and plasticity. Whether immune alterations drive schizophrenia progression is an important question to be addressed by future research, especially in light of the growing interest in applying immunotherapies in schizophrenia.

## Introduction

Schizophrenia is a severe psychiatric disease that disturbs multiple aspects of mental function. Although its etiology remains poorly understood, liability is largely genetically mediated.^1^ In the largest genome-wide association study (GWAS) yet conducted (n = 35,476 cases, 46,839 controls), over 100 independent loci were robustly associated with the disease.^2^ This GWAS represents a significant advance towards defining the molecular parts list for schizophrenia, and provides an opportunity to integrate genetic information with existing biological data to test specific etiological hypotheses.

Among many hypotheses of schizophrenia etiology, the longstanding immune theory posits that disregulation of the immune system causes schizophrenia in at least a subset of patients.^3–6^ It is known that immune components such as MHC class I,^7^ TNF-α,^8^ complement,^9^ TGF-β,^10^ and IL-6^11^ regulate brain development and adult neural plasticity. Exposure to the wrong level of an immune factor at the wrong time may consequently disrupt brain development and adult neural functioning, as supported by *in utero* immune activation in rodents^12^ and primates.^13^ Many schizophrenia patients show hallmarks of immune disease – such as prior infection,^14^ presence of autoantibodies,^15^ co-occurring autoimmunity,^16^ and inflammation,^17, 18^ – supporting the idea that immune disturbances may play a role in schizophrenia by disrupting brain development and/or adult neural function.

Given the immune disturbances apparent among schizophrenia patients, there is considerable interest in treating the disease with immune-modulating drugs.^19^ Nonspecific anti-inflammatory agents (e.g. aspirin) have shown modest efficacy among schizophrenia patients,^20^ and randomized controlled trials with more targeted immunotherapies (e.g. Tocilizumab, a monoclonal antibody against the IL-6 receptor)^21^ are currently underway. Importantly, the underlying cause of immune perturbation in schizophrenia remains unknown. A combination of genetic and environmental risk factors has been proposed to initiate immune abnormalities among schizophrenia patients.^22^ Alternatively, the immune disturbances may be epiphenomena driven by disease pathogenesis, exposure to antipsychotic drugs, or lifestyle factors associated with schizophrenia such as smoking. Immune profiling studies of schizophrenia have primarily been cross-sectional in nature, precluding causal inferences. Furthermore, important factors influencing immune status – such as hospitalization, age, sex, body mass index, diet, smoking, and medication use^23^ – are associated with schizophrenia case status^23^ and are not always accounted for. Thus, it remains unclear from the existing literature whether the relationship between schizophrenia and immune disturbances is causal, correlative, or an epiphenomenon. Adjunct immunotherapy may be a viable therapeutic option regardless of the role of immune dysregulation – whether it causes schizophrenia, influences disease progression, or is a biomarker for disease. However, clarifying the role of immune processes in schizophrenia has important implications for understanding disease biology, optimizing the timing of immunotherapy interventions, and developing effective targeted therapies.

If genetic variants influencing immune function were associated with elevated risk of schizophrenia, this would provide strong evidence that immune abnormalities are causal drivers of disease. Although genetic data have been interpreted as supporting immune involvement,^22, 24, 25^ largely due to the strong association of single-nucleotide polymorphism (SNP) alleles in the extended major histocompatibility complex (xMHC),^24^ previous studies have reported conflicting results with respect to immune pathway involvement. For instance, in gene-set enrichment analysis of the Genetic Association Information Network (GAIN) schizophrenia GWAS (1,158 cases and 1,378 controls), three of the seven overrepresented pathways were related to the immune system (TGF-β, TNFR1, and TOB1 pathways).^26^ In contrast, a more recent study integrating results across five different pathway analysis methods observed enrichment of TGF-β signaling after pooling GWAS results for major depressive disorder (9,227 cases and 7,383 controls), bipolar disorder (6,990 cases and 4,820 controls), and schizophrenia (9,379 cases and 7,736 controls) but no enrichment of immune pathways in schizophrenia alone.^27^ Thus, it remains unclear whether the immune disturbances apparent in schizophrenia are genetically mediated.

Here we directly tested the hypothesis that common variation within immune genes contributes to schizophrenia in a total sample of 35,476 schizophrenia cases and 46,839 controls. We first evaluated the collective association and overall enrichment of SNPs within immune loci in schizophrenia, to discern whether existing genetic data support an immunological cause of the disease. We then evaluated association of individual immune components with schizophrenia, to identify candidates driving the immune disturbances observed in the disease.

## Subjects and Methods

### Samples and quality control

An overview of GWAS datasets analyzed, including information about immune SNP coverage, is provided in **Table 1** and **Supplementary Table 1**. All analyses used imputed genotype dosages or summary statistics generated according to quality control and imputation protocols described in the original GWASs.

**Table 1.** Description of samples

| Sample | PMID | Cases | Controls | Cohorts |
| --- | --- | --- | --- | --- |
| Schizophrenia | 25056061 | 35,476 | 46,839 | 52 |
| Crohn's disease | 21102463 | 6,333 | 15,056 | 6 |
| Multiple sclerosis | 21833088 | 9,772 | 17,376 | 23 |
| Psoriasis | 20953190 | 2,178 | 5,175 | 1 |
| Rheumatoid arthritis | 20453842 | 5,539 | 20,169 | 6 |
| Ulcerative colitis | 21297633 | 6,687 | 19,718 | 6 |

#### Schizophrenia

We analyzed data from the most recent schizophrenia GWAS conducted by the PGC.^2^ The full dataset comprised 52 cohorts totaling 35,476 cases and 46,839 controls, and was described in detail in the primary GWAS analysis.^2^ For analyses using individual-level SNP data, we analyzed the 36 European ancestry case-control cohorts for which we had ethics approval (25,629 cases and 30,976 controls, see **Supplementary Table 2** for details). For analyses using summary statistics we used publicly available summary results generated from the complete dataset, as described in the primary analysis.^2^

#### Autoimmune diseases

To evaluate the robustness of our approach to evaluate immune enrichment, and to benchmark our findings for schizophrenia, we analyzed GWAS summary statistics from five diseases of known immune origin: Crohn’s disease (6,333 cases and 15,056 controls),^28^ multiple sclerosis (9,772 cases and 17,376 controls),^29^ psoriasis (2,178 cases and 5,175 controls),^30^ rheumatoid arthritis (5,539 cases and 20,169 controls),^31^ and ulcerative colitis (6,687 cases and 19,718 controls).^32^ Multiple sclerosis, psoriasis, and rheumatoid arthritis have long been considered classic autoimmune diseases based on the presence of self-reactive immune cells, directed against a tissue-specific antigen. Crohn’s disease and ulcerative colitis have historically been considered inflammatory diseases, but recent genetic data also support an autoimmune component.^33^ For brevity, we refer to these five immune diseases of inflammatory and autoimmune origin as *autoimmune* throughout the manuscript.

Access to the multiple sclerosis dataset^29^ was obtained with permission from the International Multiple Sclerosis Genetics Consortium (IMSGC). Access to the psoriasis GWAS dataset^30^ was obtained with permission from the Wellcome Trust Case Control Consortium (WTCCC), and imputed as described in Tsoi *et al*.^34^ The remaining autoimmune disease GWAS datasets were publicly available (see **Supplementary Table 1** and **Web Resources** for details).

### Gene sets

#### Immune gene set

We defined immune genes as those with an immune response annotation in Kyoto Encyclopedia of Genes and Genomes, Gene Ontology, Ingenuity, and Immunology Database and Analysis Portal as accessed on Sept 21, 2014 (for details, see **Supplementary Table 3**). We included autosomal genes present in three of the four databases in our immune gene set. We excluded immune genes encoded in the xMHC (chromosome 6, 25 – 34 Mb), due to the broad association and extensive LD in this region. All SNPs within the xMHC were also excluded from subsequent analyses.

Immune SNPs were defined as those falling within 50 kb upstream or downstream^2^ of the transcribed region of genes in the immune gene list (973 genes representing 145 Mb, see **Supplementary Table 1** and **Supplementary Table 4** for details of immune genes and SNP coverage).

#### Null gene set

To create the null gene set we randomly selected 973 autosomal genes outside of the xMHC (representing 150 Mb), resulting in a list containing the same number of genes as the immune gene set. Null SNPs were defined using a 50 kb gene window.

#### Brain gene set

We used the brain gene set identified and previously described by Raychaudhuri *et al*,^35^ with exclusion of those brain genes encoded within the xMHC. Briefly, brain genes were identified using four independent approaches: preferential gene expression in the brain compared to other tissues, neural-activity annotation by Panther, learning annotation in Ingenuity, and synapse annotation in Gene Ontology. Brain SNPs were defined using a 50 kb gene window (2,635 genes representing 589 Mb).

### Statistical methods

#### Association of immune genes in schizophrenia

To formally evaluate statistical significance of the immune hypothesis of schizophrenia, we performed a joint association analysis of all immune SNPs. This analysis included the 36 European ancestry case-control cohorts for which we had ethics approval (25,629 cases and 30,976 controls). Schizophrenia case status was permuted 100 times, an approach that created a null dataset while preserving the LD pattern between the 346,253 immune SNPs available for analysis. For each permutation, association testing for each immune SNP was performed by logistic regression separately in each cohort adjusting for ten multi-dimensional scaling components (C1, C2, C3, C4, C5, C6, C7, C9, C15, C18), followed by inverse-variance weighted fixed effects meta-analysis. A sum of the Wald χ^2^ (1 degree of freedom) test statistics for immune SNPs was obtained, and the empirical *p*-value was calculated as the proportion of permutation samples whose sum statistic was larger than that in the observed sample. The same permutation analysis was repeated for the null set of 290,239 SNPs within 973 randomly selected genes as a baseline comparison.

#### Benchmarking immune involvement using stratified LD Score regression

To benchmark genetically mediated immune involvement in schizophrenia, we applied stratified linkage disequilibrium (LD) Score regression, which partitions heritability into functional categories while adjusting for LD-induced correlations and accepts summary statistics as input.^36^ This method leverages the relationship between LD and association test statistics to estimate the per-SNP heritability attributable to a functional category by multiple regression of the association test statistic (χ^2^) for SNP *j* against the LD Score of SNP *j* with respect to each functional category. Briefly, the regression coefficients obtained by multiple regression correspond to category specific per-SNP heritabilities (τ_c_) that account for the effects of all other categories, and can be used to estimate category specific enrichment of SNP-based heritability (h^2^_SNP_). Thus, stratified LD Score regression identifies a functional category as enriched for heritability if SNPs with high LD to that category have higher association test statistics than SNPs with low LD to that category. The enrichment estimates are defined as the proportion of SNP heritability explained by a functional category, normalized to the proportion of SNPs in that functional category. The statistical framework has been described in detail previously.^36^

To apply stratified LD Score regression we considered only the subset of SNPs with available summary statistics that overlapped HapMap Project Phase 3 (HapMap3)^37^ SNPs (as a proxy for well-imputed SNPs), because stratified LD Score regression does not account for imperfect imputation. First, we calculated LD Scores for these HapMap3 SNPs with respect to the immune and brain SNP categories, as well as a baseline gene category that included all SNPs within a 50 kb window of the transcribed region of any gene. Next, we estimated enrichment of heritability for the immune and brain functional categories using a multiple linear regression model that included either the immune or brain annotations in addition to 54 overlapping categories (our baseline gene category, and the 53 annotations previously described by Finucane *et al*.^36^), because the accuracy of enrichment estimates is improved by including many functional categories in the model. Standard error estimates were obtained by block jackknifing over 200 equally-sized blocks of SNPs.^36^

#### Immune candidate gene identification

To evaluate association of specific immune components with schizophrenia, we used summary statistics from the PGC schizophrenia GWAS.^2^ Index SNPs, defined as the SNP with the smallest association *p*-value for each disease-associated locus, were identified in the primary analysis of the PGC dataset as previously described.^2^

## Results

### Immune gene set and corresponding immune SNPs

To define an inclusive immune gene set we used an annotation-based approach which captured 973 autosomal immune genes, represented by a total of 587,933 SNPs in the schizophrenia GWAS (see **Supplementary Table 4** for details of SNP coverage for immune genes). Although there is a robust xMHC association in schizophrenia, there is extensive LD in this region which can bias standard enrichment approaches and lead to false-positive results^38^. To avoid this bias, we excluded the xMHC from the main analyses, and focused on immune genes outside of this region. To substantiate that our approach did not exclude from investigation immune loci with a major role in schizophrenia, we fine-mapped the xMHC association in schizophrenia (**Supplementary Methods**). Concordant with previous reports,^39, 40^ the primary signals in schizophrenia captured by GWAS variants did not map to coding variants in the human leukocyte antigen (HLA) genes typically implicated in autoimmune disease. Instead, we found three independent xMHC associations in schizophrenia corresponding to SNPs that represented 1) an extended class I region of association spanning ~2 Mb, 2) a class II region variant located near the C4 gene, concordant with the recent finding that C4 structural variants – requiring specialized imputation methods beyond standard GWAS analysis pipelines – are associated with schizophrenia^40^, and 3) the *SYNGAP1* gene, in which *de novo* mutations have already been implicated in schizophrenia^41^ and other neurodevelopmental disorders^42, 43^ (**Supplementary Figure 1**). Although we did not impute C4 structural variants in the present study, Sekar *et al*. have previously reported that there is no remaining association in the class II region after adjusting for C4 variation in the PGC schizophrenia GWAS.^40^ Thus, despite previous interpretations of the xMHC association in schizophrenia representing an autoimmune cause of disease^22, 24, 25^, we found no evidence for involvement of the HLA genes typically driving autoimmune susceptibility. We cannot disprove that genetic variation in the MHC may influence schizophrenia susceptibility via additional independent variants that did not reach significance in the present analysis, or underlying causal variants that were not captured in the current GWAS. Nevertheless, our findings suggest that our focus on immune genes outside of the xMHC was unlikely to have excluded from investigation common immune variants captured in the current GWAS that have a major role in schizophrenia.

### Evaluating collective association of immune genes in schizophrenia

To determine whether current genetic data support an immune cause of schizophrenia, we first evaluated the summed association signal from genes encoding immune components using individual-level SNP data from the largest GWAS currently available^2^ (25,629 cases and 30,976 controls of European ancestry). We observed evidence of inflation of the summed association test statistics for immune loci in schizophrenia (λ_immune_ = 1.48, empirical *p* < 0.01, **Figure 1**), suggesting potential involvement of immune pathways in disease etiology. Given the substantial polygenic contribution to schizophrenia, some degree of inflation is expected even among a randomly selected set of SNPs.^2^ To determine whether the collective association of immune SNPs exceeded that expected given the polygenic architecture of schizophrenia, we repeated the permutation analysis on a set of 973 randomly selected autosomal genes representing approximately the same proportion of SNPs as our immune gene set. We observed greater inflation of the summed association test statistic for this null gene set (λ_null_ = 1.54, empirical p < 0.01, **Figure 1**), suggesting the collective association observed for immune SNPs was driven by the polygenicity of schizophrenia rather than specific involvement of immune loci in the disease.

**Figure 1.**
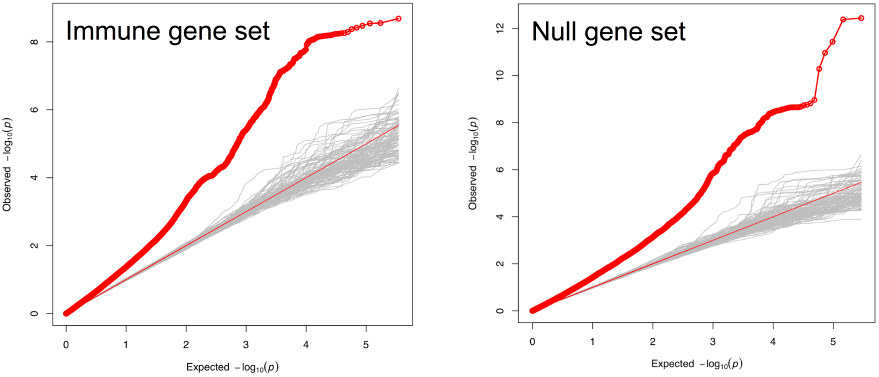
Evaluation of the immune hypothesis in schizophrenia. Quantile-quantile plots of 346,253 SNPs representing 973 immune genes (left) and 209,239 SNPs representing 973 randomly selected genes (right). Association testing was done in the 36 European ancestry case-control cohorts for which individual-level genotype data was available (25,629 cases and 30,976 controls). Observed association statistics (red) and those from 100 phenotype-permuted replicates (gray) are shown.

### Benchmarking contribution of immune genes to schizophrenia

To benchmark genetically mediated immune pathway involvement, we quantified enrichment of heritability among immune SNPs compared to SNPs in the rest of the genome in schizophrenia and five diseases of known immune origin (Crohn’s disease,^28^ multiple sclerosis,^29^ psoriasis,^30^ rheumatoid arthritis,^31^ and ulcerative colitis^32^). We estimated enrichment of immune SNPs using the recently developed stratified LD Score regression approach,^36^ which uses multiple linear regression of χ^2^ test statistics against LD Score with respect to functional categories to estimate category specific enrichment of SNP-based heritability (h^2^_SNP_). As expected based on previous literature and known biology, immune genes were consistently enriched 2- to 8-fold for h^2^_SNP_ across all five autoimmune diseases (p < 5×10^−3^, **Figure 2** and **Table 2**). In contrast to our findings in autoimmune diseases, immune genes were not enriched for heritability in schizophrenia (p = 0.94, **Figure 2** and **Table 2**). As a separate approach we applied stratified false discovery rate (sFDR) control^44^ to obtain enrichment estimates for immune genes, and again observed immune enrichment across the five autoimmune diseases but not schizophrenia (**Supplementary Figure 2** and **Supplementary Methods** in the Supplement). Taken together, these results suggest that immune genes as a group may not be major drivers of schizophrenia risk.

**Figure 2.**
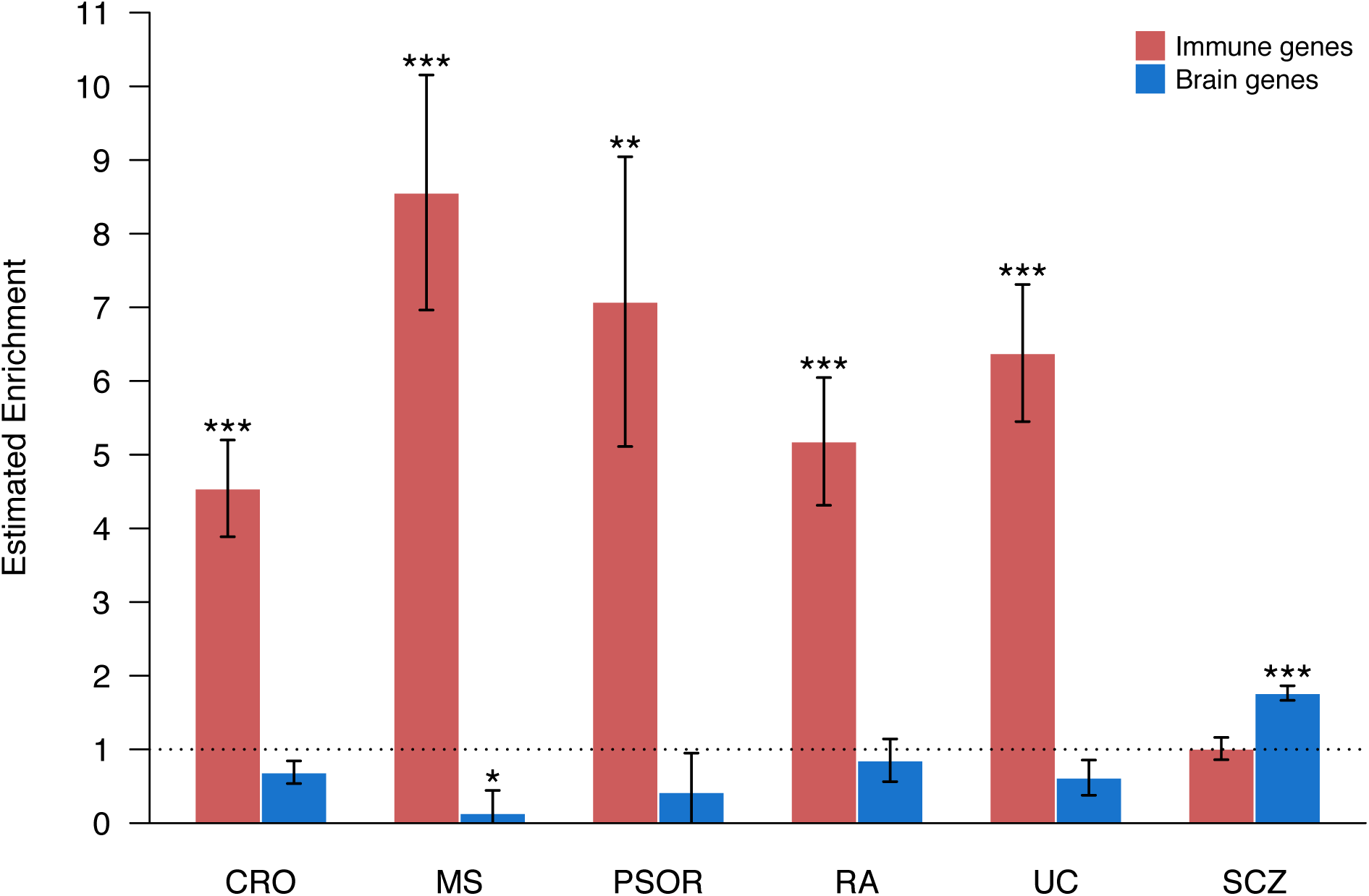
Estimated enrichment for immune (red) and brain (blue) genes in schizophrenia and five autoimmune diseases. The *y*-axis represents estimated enrichment for each gene set, defined as the proportion of heritability explained divided by the proportion of SNPs for that gene set. Values > 1 (dotted line) indicate enrichment of heritability. Error bars represent enrichment estimates ± standard error. ***p < 1×10^−5^, ** < 1×10^−3^, *p < 0.01 in a test of whether the estimated enrichment was equal to one. CRO, Crohn’s disease; MS, multiple sclerosis; PSO, psoriasis; RA, rheumatoid arthritis; UC, ulcerative colitis; SCZ, schizophrenia.

**Table 2.** Enrichment and per-SNP heritability estimates for immune and brain gene sets

| Disease | $h^2$ enrichment | SE | p | per-SNP $h^2$ ($\tau_c$) | SE | p |
| --- | --- | --- | --- | --- | --- | --- |
| Immune gene set |  |  |  |  |  |  |
| Schizophrenia | 1.01 | 0.15 | 0.94 | $-7.01 \times 10^{-9}$ | $1.11 \times 10^{-8}$ | 0.53 |
| **Crohn's disease** | **4.54** | **0.66** | **$6.80 \times 10^{-8}$** | **$2.04 \times 10^{-7}$** | **$6.28 \times 10^{-8}$** | **$1.16 \times 10^{-3}$** |
| **Multiple sclerosis** | **8.56** | **1.60** | **$2.16 \times 10^{-6}$** | **$1.49 \times 10^{-7}$** | **$2.84 \times 10^{-8}$** | **$1.55 \times 10^{-7}$** |
| **Psoriasis** | **7.08** | **1.97** | **$2.00 \times 10^{-3}$** | **$2.26 \times 10^{-7}$** | **$7.17 \times 10^{-8}$** | **$1.62 \times 10^{-3}$** |
| **Rheumatoid arthritis** | **5.18** | **0.87** | **$1.41 \times 10^{-6}$** | **$1.01 \times 10^{-7}$** | **$2.70 \times 10^{-8}$** | **$1.83 \times 10^{-4}$** |
| **Ulcerative colitis** | **6.38** | **0.93** | **$7.36 \times 10^{-9}$** | **$1.87 \times 10^{-7}$** | **$3.68 \times 10^{-8}$** | **$3.74 \times 10^{-7}$** |
| Brain gene set |  |  |  |  |  |  |
| **Schizophrenia** | **1.76** | **0.10** | **$1.14 \times 10^{-14}$** | **$4.04 \times 10^{-8}$** | **$9.24 \times 10^{-9}$** | **$1.23 \times 10^{-5}$** |
| Crohn's disease | 0.69 | 0.15 | 0.04 | $-1.54 \times 10^{-8}$ | $2.20 \times 10^{-8}$ | 0.48 |
| Multiple sclerosis | 0.14 | 0.31 | $4.84 \times 10^{-3}$ | $-1.27 \times 10^{-8}$ | $1.11 \times 10^{-8}$ | 0.25 |
| Psoriasis | 0.42 | 0.53 | 0.27 | $-6.30 \times 10^{-9}$ | $3.29 \times 10^{-8}$ | 0.85 |
| Rheumatoid arthritis | 0.85 | 0.29 | 0.61 | $-4.61 \times 10^{-10}$ | $1.21 \times 10^{-8}$ | 0.97 |
| Ulcerative colitis | 0.62 | 0.24 | 0.11 | $-2.49 \times 10^{-9}$ | $1.32 \times 10^{-8}$ | 0.85 |

Bold font indicates those SNP sets that are enriched based on both enrichment estimates and per-SNP $h^2$ estimates ( $\tau_c$ ). $\tau_c$ estimates are obtained using a multivariate model that includes all other functional categories whereas the enrichment estimates only consider the functional category of interest. Therefore, when the enrichment estimate is significant without a significant $\tau_c$ estimate, this suggests the result may be driven by correlation with other functional categories.

It is possible that we were unable to detect true immune enrichment in schizophrenia due to its unique genetic and clinical architecture (highly polygenic and clinically heterogeneous) relative to the autoimmune diseases analyzed. As a positive control, we applied stratified LD Score regression to quantify enrichment of a set of brain genes previously reported to be enriched for schizophrenia heritability.^45^ As expected, we observed significant enrichment of brain genes in schizophrenia (p = 1.14×10^−14^, **Figure 2** and **Table 2**), indicating that we are able to detect true pathway enrichment in schizophrenia despite the high degree of polygenicity and clinical heterogeneity.

### Identification of immune candidates individually associated with schizophrenia

Although we found no overall enrichment of immune loci in schizophrenia, we hypothesized that individual immune genes may be implicated in the disease. Of the 108 previously reported loci associated with schizophrenia,^2^ we identified six independent regions on chromosomes 2, 5, 8, 11, and 16 where the index SNP was an immune SNP (**Table 3**, **Supplementary Figure 3**). To the best of our knowledge none of these loci, which represented the *DPP4, HSPD1, EGR1, CLU, ESAM*, and *NFATC3* genes, have been previously associated with an autoimmune disease. All six of the immune genes associated with schizophrenia are expressed in human brain tissue^46^ (**Supplementary Figure 4**). Interestingly, their protein products have roles in immune cell activation and adhesion, as well as established roles in the brain such as regulating myelination (*HSPD1*),^47^ synaptic plasticity (*EGR1*),^48^ blood-brain-barrier permeability (*ESAM*),^49^ and neuronal loss after brain injury (*NFATC3*,^50^ *CLU*^51^).

**Table 3.** Genome-wide significant immune genes in schizophrenia

| SNP | Chr | Position | OR (95% CI) | *p* | Gene | Location<sup>a</sup> |
| --- | --- | --- | --- | --- | --- | --- |
| rs2909457 | 2 | 162845855 | 0.94 (0.92 - 0.96) | 4.38e-08 | *DPP4* | + 2.9 kb |
| rs6434928 | 2 | 198304577 | 0.93 (0.91 - 0.95) | 1.48e-11 | *HSPD1* | + 46.7 kb |
| rs3849046 | 5 | 137851192 | 1.06 (1.04 - 1.08) | 4.83e-09 | *EGR1* | Intronic |
| rs73229090 | 8 | 27442127 | 0.91 (0.88 - 0.94) | 1.95e-08 | *CLU* | + 12.3 kb |
| rs55661361 | 11 | 124613957 | 0.92 (0.90 - 0.94) | 3.68e-12 | *ESAM* | + 9.1 kb |
| rs8044995 | 16 | 68189340 | 1.08 (1.05 - 1.11) | 3.27e-08 | *NFATC3* | Intronic |

<sup>a</sup>Location relative to immune gene of interest; +, downstream.

## Discussion

The hypothesis that schizophrenia may be an immunological disease is longstanding.^3–5^ Using the largest collection of genetic data currently available, we evaluated the immune hypothesis of schizophrenia empirically. We have shown that common variation at immune genes presents a very different genetic architecture in schizophrenia as compared to diseases of known immune origin. First, the collective association of immune genes outside of the MHC in schizophrenia was not greater than expected given the polygenic architecture of the disease. Second, there was no enrichment of heritability among non-MHC immune genes in schizophrenia, in contrast to that observed in autoimmune diseases. While broad immune enrichment – or enrichment of specific immune pathways such as TGF-β signaling, which has already been observed when pooling GWAS results across all major adult psychiatric disorders^27^ – may be detected as GWAS sample sizes increase further, our benchmarking establishes that the degree of enrichment in schizophrenia is substantially less than that seen in autoimmune diseases. Third, the immune loci that were individually associated with schizophrenia had important alternate roles in brain development and homeostasis, raising the possibility that proteins with dual immune-neural function are responsible for the link between schizophrenia and the immune system. For instance, ESAM regulates blood-brain-barrier permeability,^49^ and may influence susceptibility to schizophrenia by regulating exposure to the peripheral immune milieu. As the sample sizes available for genetic study increase and new genotyping approaches emerge, additional immune genes robustly associated with schizophrenia are likely to be discovered, and it will be critical to evaluate how these immune components may act in the brain.

Our methodological approach was subject to six potentially important limitations. Firstly, the immune gene list was broad, which may have diluted any enrichment in a more specific immune subset or pathway. Secondly, our annotation-based approach to defining the immune gene list did not capture remote regulatory regions for immune genes. Thirdly, the MHC region was excluded from all analyses subsequent to fine-mapping the MHC association. Due to the extensive LD in the MHC region, current approaches for gene-enrichment analysis are not robust to inclusion of MHC variants and this is an important area of future methods development. Fourthly, schizophrenia is an umbrella diagnosis that, like other complex disorders, likely captures many distinct molecular subtypes.^52^ Thus, the broad phenotype classification used to define the patient cohort in the present study likely resulted in clinical heterogeneity within our sample. As immune disturbances may be causal in only some subset of schizophrenia patients, this clinical heterogeneity may have diluted association signals in the immune gene set. Fifthly, our study was limited to common variants captured in current GWASs, which do not account completely for the estimated heritability of schizophrenia.^53^ Finally, given evidence that exposure to inflammatory mediators *in utero* increases the risk of schizophrenia,^54^ it may be genetic variation in the mother’s immune loci that contributes to the immune disturbances seen in patients with schizophrenia. Given these limitations, we cannot completely exclude a potential genetic etiology for the immune disturbances observed in schizophrenia.

Despite these limitations, to the best of our knowledge this was the most comprehensive investigation of the hypothesis that immune genes contribute to schizophrenia. Based on current GWAS data schizophrenia does not appear to be an autoimmune disease *perse*, although there may be modest contributions to genetic susceptibility from a specific subset of immune genes with additional roles outside of immunity (for example, neurodevelopmental). Our findings also raise the possibility that the immune disturbances observed in schizophrenia are of non-genetic etiology. Importantly, we cannot exclude the possible causal role of environmental risk factors that activate or “prime” the immune response (e.g. infections, stress) in schizophrenia. Alternatively, the immune disturbances seen in schizophrenia may be a downstream factor in disease pathogenesis, fuelling progression or modifying disease outcomes rather than initiating the disease. Finally, the immune changes observed in schizophrenia may simply be a by-product of disease pathogenesis or patient lifestyle factors (i.e. antipsychotic medication, smoking, and diet). Whether the immune abnormalities accompanying schizophrenia are causal, disease modifying, or epiphenomena is an important question to be addressed by longitudinal studies, particularly given burgeoning interest in potential immunotherapies.

## Web Resources

- Summary statistics:

- Schizophrenia:^2^ http://www.med.unc.edu/pgc/files/resultfiles/scz2.snp.results.txt.gz%20%20
- Crohn’s disease:^28^ ftp://ftp.sanger.ac.uk/pub4/ibdgenetics/cd-meta.txt.gz
- Rheumatoid arthritis:^31^ http://www.broadinstitute.org/ftp/pub/rheumatoid_arthritis/Stahl_etal_2010NG/
- Ulcerative colitis:^32^ ftp://ftp.sanger.ac.uk/pub4/ibdgenetics/ucmeta-sumstats.txt.gz
- Stratified LD Score regression software:^36^ github.com/bulik/ldsc
- sFDR software:^44^ http://www.utstat.toronto.edu/sun/Software/SFDR

## Acknowledgements

### Acknowledgement of Funding Sources

VF Goncalves is supported by CIHR operating grant MOP 115097. S Raychaudhuri is supported by NIH grants U19-AI111224-01 and U01-HG007033-03. JL Kennedy is supported by CIHR grant MOP 115097. J Knight is the Joanne Murphy Professor in Behavioural Science. JG Pouget is supported by Fulbright Canada, the Weston Foundation, and by Brain Canada through the Canada Brain Research Fund, a public-private partnership established by the Government of Canada. The funding sources did not influence the study design, data analysis, or writing of this manuscript.

### Additional Acknowledgements

Computations were performed on the CAMH Specialized Computing Cluster (SCC), which is funded by The Canada Foundation for Innovation, Research Hospital Fund. We also thank SURFsara (www.surfsara.nl) for the support in using the Lisa Compute Cluster. We thank Richard Trembath, PhD (Queen Mary University of London) and Jonathan Barker, MD (King’s College London) for allowing us access to the imputed psoriasis GWAS data, and the IMSGC for providing us with access to the multiple sclerosis GWAS summary data. We also thank Lei Sun, PhD (University of Toronto) for providing the sFDR scripts, Lisa Strug, PhD (University of Toronto) for feedback on study design and the interpretation of the sFDR results, and Kamil Slowikowski, BSc (Harvard University) and Harm-Jan Westra, PhD (Harvard University) for assistance writing and troubleshooting scripts for the permutation analysis.

